# TP53 mutations as potential prognostic markers for specific cancers: Analysis of data from The Cancer Genome Atlas and the International Agency for Research on Cancer TP53 Database

**DOI:** 10.1101/368027

**Authors:** Victor Li, Karen Li, John T. Li

## Abstract

Mutations in the tumor suppressor gene TP53 are associated with a variety of cancers. Therefore, it is important to know the occurrence and the prognostic effects of TP53 mutations in certain cancers. Over 29,000 cases of TP53 mutations were obtained from the April 2016 release of the Internal Agency for Research on Cancer (IARC) TP53 Database, and 7,893 cancer cases were compiled in the cBioPortal for Cancer Genomics from the 33 most recent studies of The Cancer Genome Atlas (TCGA). The data was analyzed, and it was found that the majority of TP53 mutations were missense and the major mutational hotspots were located at codons 248, 273, 175, and 245 in exons 4–8 for somatic mutations with the addition of codon 337 and other mutations in exons 9–10 for germline mutations. Out of 33 TGCA studies, the effects of TP53 mutations were statistically significant in ten cancers (ovarian serous cystadenocarcinoma, lung adenocarcinoma, hepatocellular carcinoma, head and neck squamous cell carcinoma, acute myeloid leukemia, clear cell renal cell carcinoma (RCC), papillary RCC, chromophobe RCC, uterine endometrial carcinoma, and thymoma) for survival time and in six cancers (ovarian serous cystadenocarcinoma, pancreatic adenocarcinoma, hepatocellular carcinoma, chromophobe RCC, acute myeloid leukemia, and thymoma) for disease-free survival time. Also, it was found that the most common p53 mutation in hepatocellular carcinomas (R249S) was a much better indicator for poor prognosis than TP53 mutations as a whole.

**Author summary:** The TP53 gene codes for the tumor suppressor protein, p53, which is essential for DNA repair, cell cycle arrest, and apoptosis. It is commonly inactivated or partially disabled by mutations, contributing to the development of a variety of human cancers. In this study, over 29,000 cases from the April 2016 release of the International Agency for Research on Cancer TP53 Database (IARC) were analyzed to determine the distribution of mutations in the TP53 gene. Data was also collected from the 33 most recent The Cancer Genome Atlas (TCGA) studies to determine the prevalence of TP53 mutations in cancers and their effects on survival and disease-free survival time. It was found that there were statistically significant differences between cases with and without TP53 mutations in ten cancers when comparing survival time, and in six cancers when comparing disease-free survival time. This indicates that TP53 mutations are potential prognostic markers that can be used to further improve the accuracy of predicting the survival time and disease-free survival time of cancer patients.

## Introduction

The tumor suppressor gene TP53 encodes for the p53 protein, which serves a role in DNA repair, cell cycle arrest, apoptosis, and other pathways that prevent the development of cancers. The TP53 gene and its corresponding protein are frequently inactivated or partially disabled by mutations that lead to increased risks of developing cancer. Somatic TP53 mutations are very frequent in most human cancers, occurring in 5 to 80% of them, depending on the cancer type and stage [1].

### Domains of p53

The TP53 gene contains 19,200 nucleotides in its 11 exons and 10 introns. Of the 11 exons, exon one, the first half of exon two, and the majority of exon 11 are non-coding exons [2]. The two transactivation domains are encoded in exons 2–4, the DNA-binding domain in exons 4–8, the tetramerization domain in exons 9–10, and the basic domain in exons 10–11. Each domain of this 393 amino acid protein has its own distinct function.

Transactivation domain one (AD1) and transactivation domain two (AD2) are comprised of amino acids 1–92. AD1 is required for cell cycle arrest but is dispensable for apoptosis [3]. AD2 is responsible for apoptosis and is aided by the proline-rich domain which is now part of AD2 [4, 5]. The DNA-binding domain (DBD), comprised of amino acids 102–292, is essential for the role of p53 as a sequence-specific transcription factor. The DBD of p53 binds to a specific DNA sequence to activate transcription, mediate apoptosis, and conduct cell cycle arrest to suppress the growth of tumor cells [6]. The DBD is followed by the tetramerization domain (TD), comprised of amino acids 326–356, which aids the DBD in binding to DNA and other proteins by increasing the strength of interactions between p53 and other structures [3] (Fig 1).

**Fig 1.**
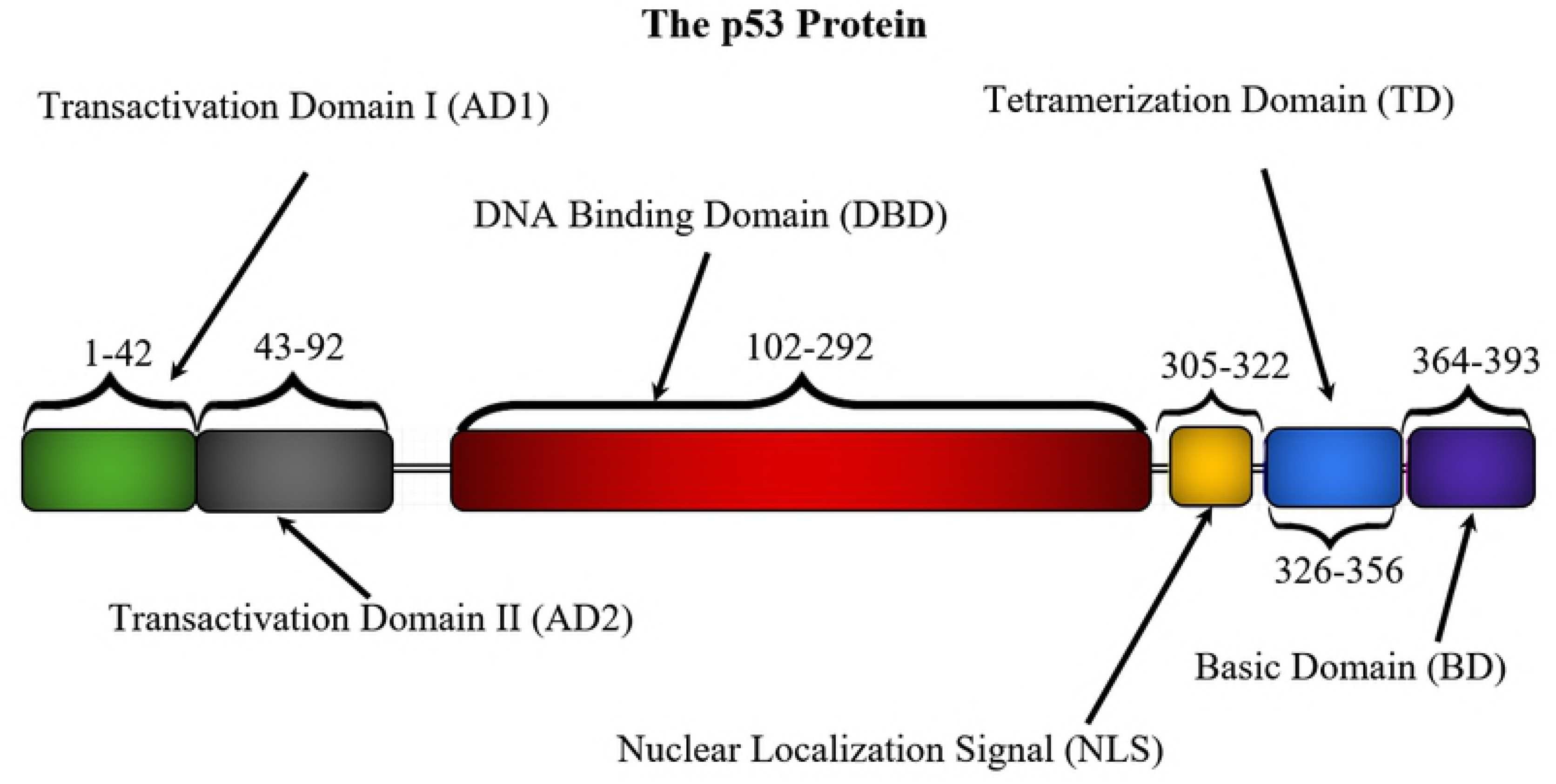
Domains of the p53 protein according to amino acid number. Data from “The Functional Domains in p53 Family Proteins Exhibit both Common and Distinct Properties.”

### Li-Fraumeni and Li-Fraumeni-like syndromes

TP53 is the only gene so far identified in which mutations are definitively associated with Li-Fraumeni (LFS) and Li-Fraumeni-like (LFL) syndromes, which predispose patients to certain types of cancers. Over 50% of families with LFS have an inherited mutation in the TP53 gene [7]. The most common germline mutations in tumors are located at amino acids 248, 337, 273, and 175 [2]. It is known that these mutations can inactivate or disrupt the function of the p53 protein and increase risks of early onset cancer [8].

### Objective of study

Past research has determined the most common TP53 mutations, examined the development of p53 alterations, and identified cancers that TP53 mutations were prevalent in. This study attempted to establish TP53 mutations as potential prognostic markers for specific cancers by investigating if the presence of TP53 mutations in certain cancers was beneficial or detrimental to survival time and disease-free survival time. The distribution of TP53 mutations at specific exons/introns and codons was examined, and the potential of using specific mutations as cancer prognostic markers was evaluated.

## Results

### Somatic mutations

The percentage of certain types of somatic TP53 mutations is unlike those of other tumor suppressors. Mutations in TP53 are 73.16% missense, 9.06% frameshift, 8.17% nonsense, 3.62% silent, 2.4% splice, 0.74% intronic, 0.17% large deletion, and 2.28% other [2] (Fig 2). Most tumor suppressors are inactivated by frameshift or nonsense mutations. However, due to the low core thermodynamic stability of p53, the protein is most often inactivated by missense mutations in TP53 that cause single amino acid changes [11, 12].

**Fig 2.**
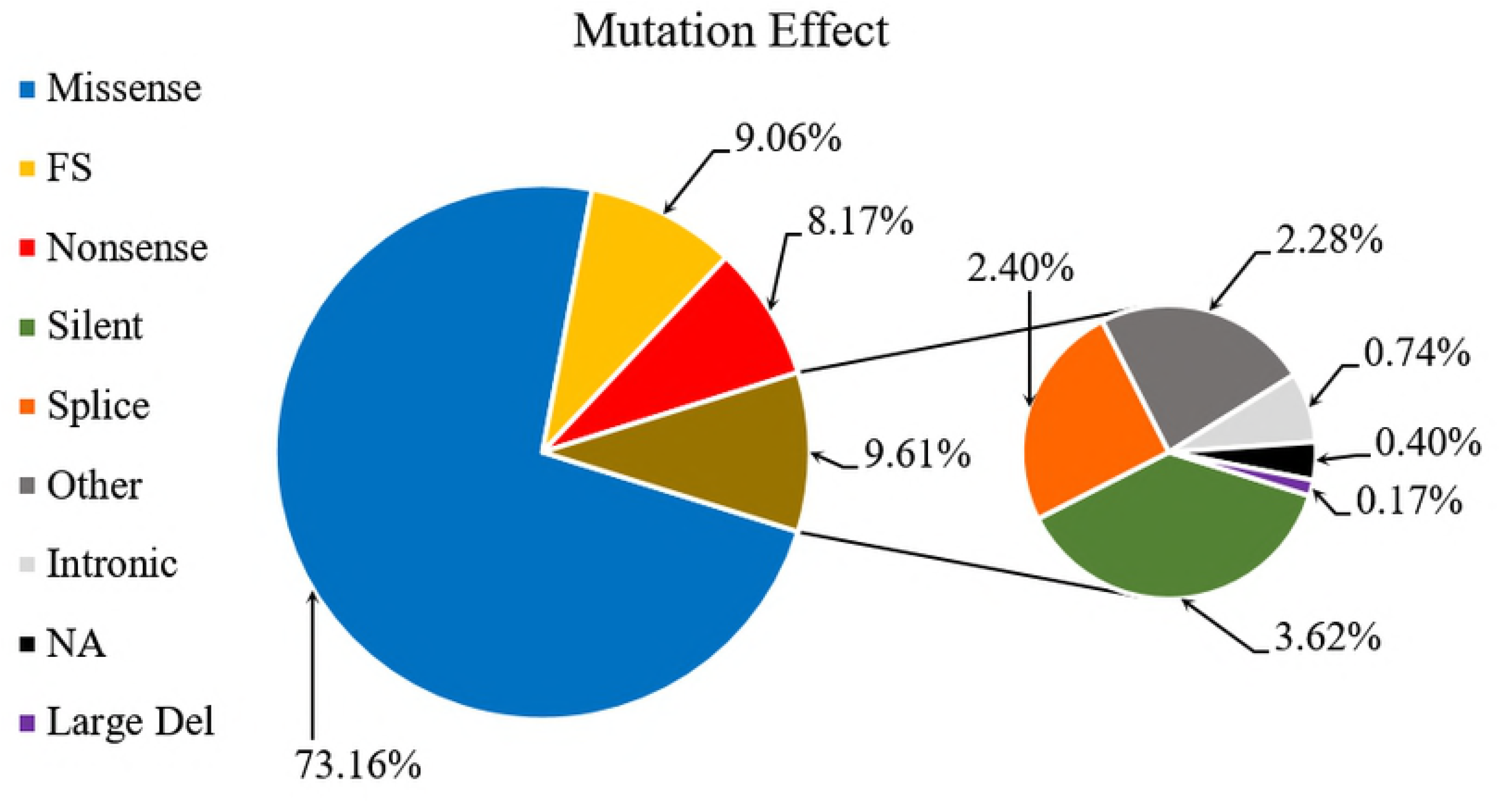
Mutation effect of TP53 somatic mutations. Pie chart shows the proportion of different types of somatic TP53 mutations. FS: frameshift; NA: not available. In total, there were 28,869 cases with mutations. Data from IARC TP53 Database (R18, April 2016).

Very few somatic mutations occur in AD1 and AD2. Even though the two transactivation domains represent 24% of the length of the gene, only 4% of somatic TP53 mutations occur in AD1 and AD2. The majority of mutations occur in the DBD, which is about the same length as AD1 and AD2. The DBD represents 26% of the TP53 gene, but almost 90% of somatic TP53 mutations occur here. 96% of somatic missense mutations occur in the DBD and 92% of somatic mutations in the DBD are missense. Only about 3% of somatic mutations are located in the TD (Fig 3A). Mutational hotspots for somatic mutations are located at codons 248, 273, 175, and 245 (Fig 4A).

**Fig 3.**
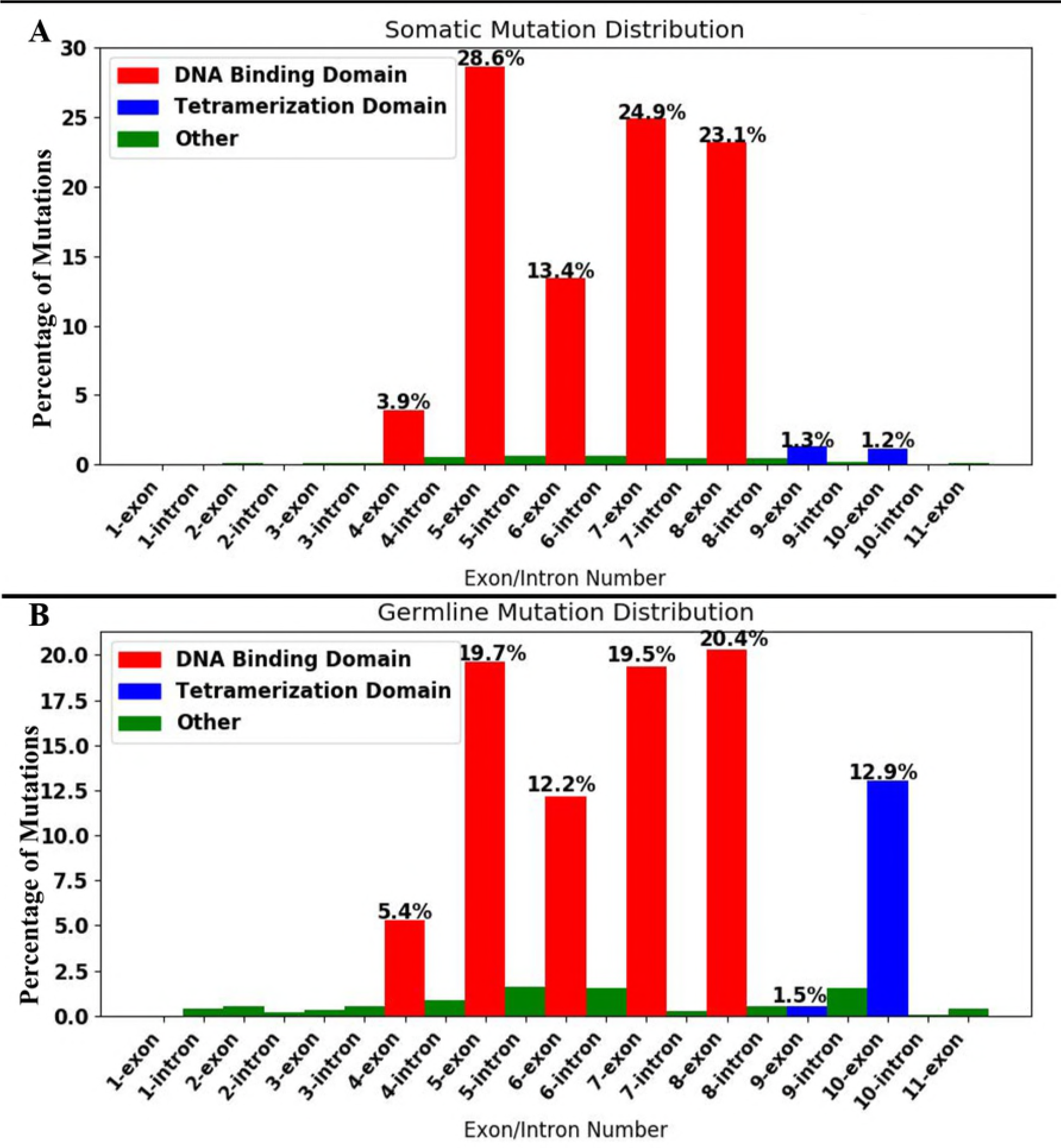
Intron/exon distributions of somatic and germline mutations. Histograms showing the percentage of mutations in specific exons and introns. Data from IARC TP53 Database (R18, April 2016).

**Fig 4.**
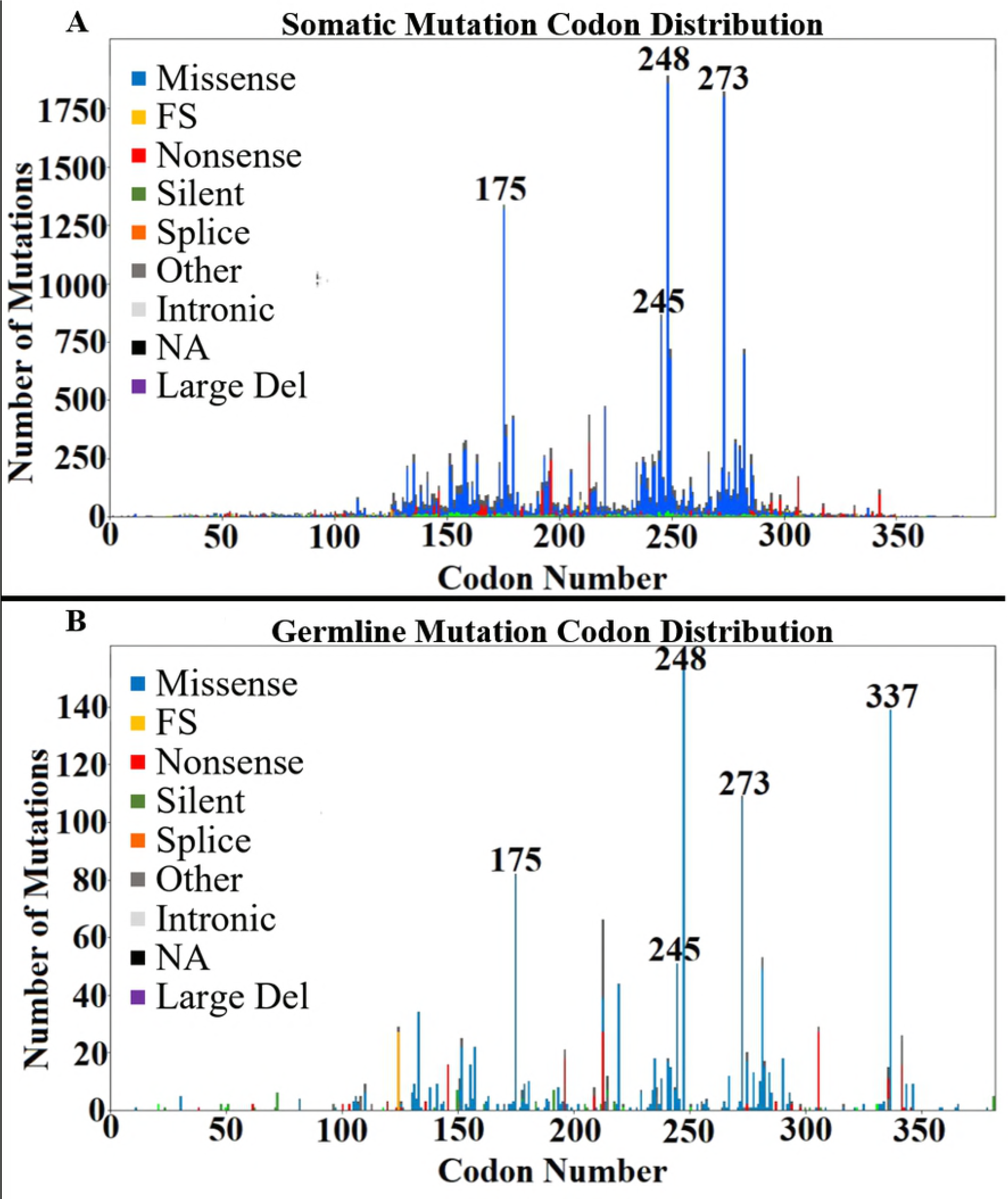
Codon distributions of somatic and germline mutations. Histogram displaying the frequency and types of mutations at certain codons. FS: frameshift; NA: not available. The major mutational hotspots are labeled. Data from IARC TP53 Database (R18, April 2016).

### Germline mutations

The distribution of germline mutations is different from the distribution of somatic mutations. 94% of all somatic TP53 mutations are located in exons 4–8, whereas only 77% of all germline TP53 mutations are located in exons 4–8. In comparison to somatic mutations, the proportion of germline mutations that occur in the DBD is lower, and the proportion of mutations in the introns, and exons of TD, AD1, and AD2 are relatively higher. The percentage of mutations in the TD is approximately 3% in somatic mutations, and approximately 14% in germline mutations. The only section of the DBD that increased in mutation proportion from somatic mutations to germline mutations was exon 4 (Fig 3B). The top three mutational hotspots occur at codons 248, 337, and 273 (Fig 4B). All three of these mutational hotspots are associated with LFS or LFL, and about 62% of the 1644 cancer cases with germline mutations had LFS, LFL, or met the TP53 Chompret criteria.

TP53 mutations are common in many different types of cancers. According to “Integrated Genomic Analyses of Ovarian Carcinoma” and “Comprehensive Genomic Characterization of Squamous Cell Lung Cancers,” two TCGA papers published in Nature, TP53 mutations occur in 96% of ovarian serous cystadenocarcinomas and 94% of lung squamous cell carcinomas [13, 14]. In this study, almost every sample of the two aforementioned cancers had TP53 mutations, but for other types of cancers such as uveal melanomas, which had a sample size of 80, there were no occurrences of TP53 mutations (Table 1).

**Table 1.**
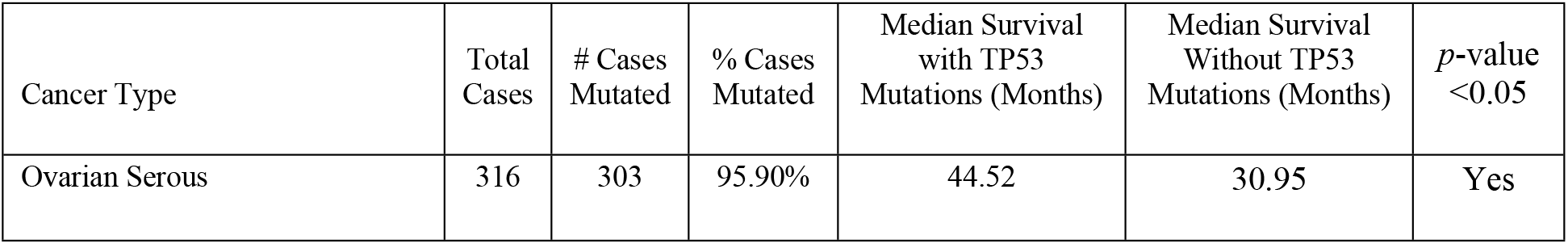
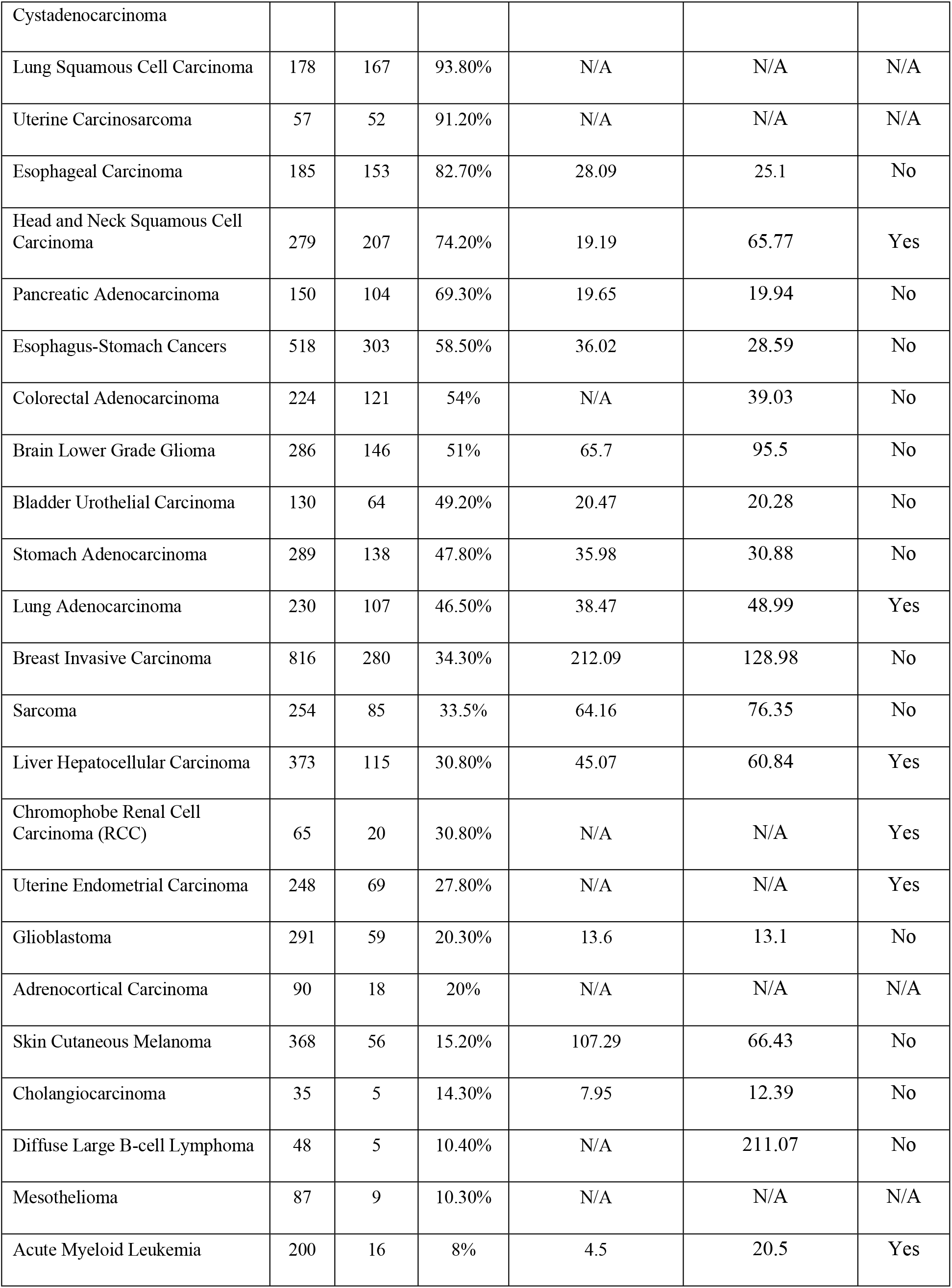
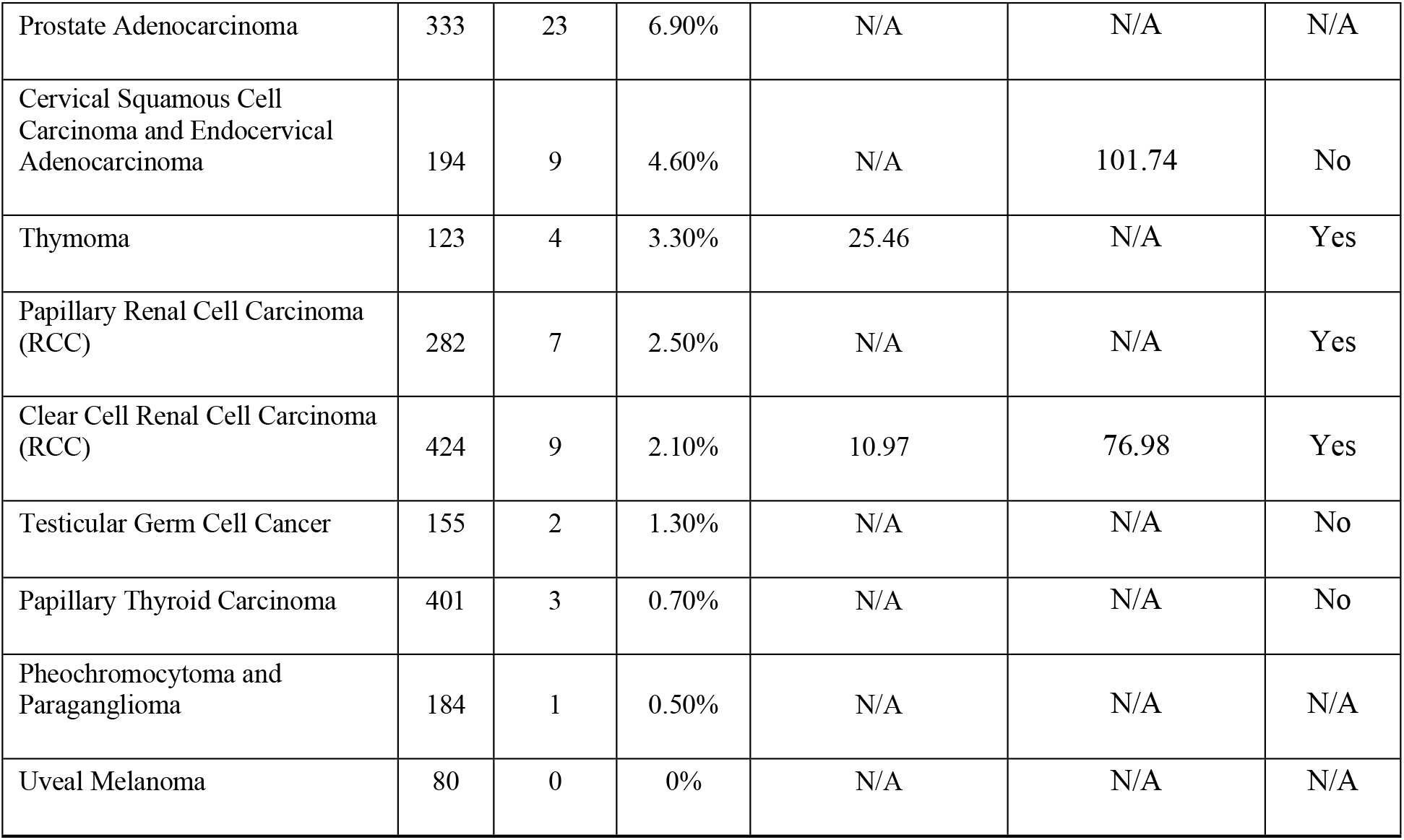
Percentage of TP53 mutations and median survival time of 33 cancers. Data from TCGA studies compiled in cBioPortal for Cancer Genomics.

### Overall survival

Of the 33 cancers analyzed, the difference in survival time of 10 cancers was found to be
statistically significant when comparing samples with and without TP53 mutations (Table 1). The cancers were separated into four groups. The first group consisted of cancers in which TP53 mutations were beneficial. The second group included cancers in which cases without TP53 mutations survived up to two times longer than cases with TP53 mutations. The third group consisted of cancers in which cases without TP53 mutations survived over three times longer than cases with TP53 mutations. Lastly, the fourth group consisted of cancers in which median survival time could not be calculated for either cases with or without TP53 mutations since >50% of patients were still alive by the end of the study.

According to previous research, TP53 mutations correlate with bad prognosis. However, in ovarian serous cystadenocarcinoma, the only cancer in group one, TP53 mutations were correlated with an increase in survival time. Out of 316 cases, 95.9% contained TP53 mutations. The median survival time was 44.52 months for cases with mutations and 30.95 months for cases without mutations. Ovarian serous cystadenocarcinoma was the only cancer type where TP53 mutations were found to be beneficial to the patient, as those with mutations survived 1.44 times longer than their wild-type counterparts (Fig 5A).

**Fig 5.**
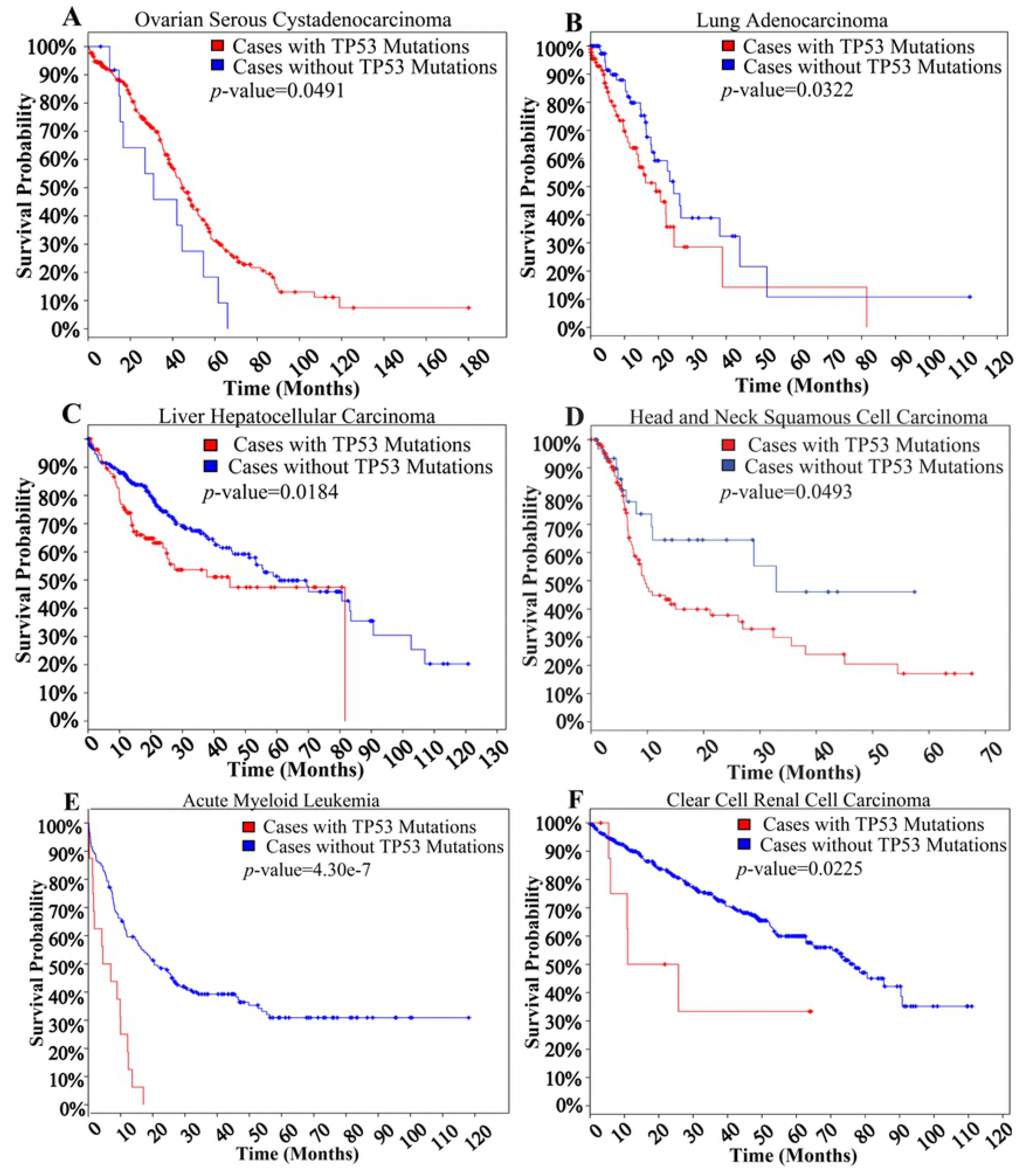
Survival Kaplan-Meier estimates of select cancers. Data from TCGA studies compiled in cBioPortal for Cancer Genomics.

The next two cancers were in the second group. The first was lung adenocarcinoma, where about 46.5% out of 230 cases had TP53 mutations. According to median survival data, cases without mutations survived for 48.9 months and cases with mutations survived for 38.47 months. Those without mutations survived 1.27 times longer. Even though mutations resulted in poor prognosis, only the minority of cases were affected negatively (Fig 5B). The second cancer was liver hepatocellular carcinoma, where 30.8% out of 373 cases had TP53 mutations. Once again, like in lung adenocarcinomas, the minority of patients were affected negatively. The median survival time was 60.84 months for cases without mutations, and 45.07 months for cases with mutations. Those without mutations survived 1.35 times longer. While it seems like the survival percentage in months 70–80 were equal between those with and without mutations, this occurred because of censoring, which resulted in no points of interest between months 50–80 for samples with TP53 mutations (Fig 5C).

The next three cancers were in the third group. The first was head and neck squamous cell carcinoma, where TP53 mutations occurred in 74.2% out of 279 cases. Median survival time was 65.77 months for cases without mutations, and 19.19 months for cases with mutations. Those without mutations survived 3.43 times longer. Out of all cancers with statistically significant differences in survival time, this was the only cancer in which the majority of patients possessed detrimental TP53 mutations (Fig 5D). The second cancer was acute myeloid leukemia, where 8.0% out of 200 cases contained TP53 mutations. By the first 10 months only 25% of cases with mutations were still alive, and by 17 months all such cases were deceased. However, for those without mutations, about 66% of cases were alive by 10 months, and by 17 months over 50% of cases were still alive. Median survival time were 20.5 months for cases without mutations, and 4.5 months for cases with mutations. Those without mutations survived 4.56 times longer. Since only 8.0% of cases had TP53 mutations, only the minority of patients were affected negatively (Fig 5E). The third was clear cell renal cell carcinoma (RCC). Although only 2.1% out of 424 cases had TP53 mutations, the most drastic differences in survival time was seen. Median survival time was 77 months for cases without mutations, and 11 months for cases with mutations. Those without mutations survived 7.02 times longer (Fig 5F).

The next four cancers were in group four. The first was papillary RCC. Out of 282 samples, only 2.5% were mutated. According to the Kaplan-Meier estimate, the upper quartile survival time was 8.48 months for cases with TP53 mutations and 67.12 months for cases without TP53 mutations. The second was chromophobe RCC, which had TP53 mutations in 30.8% of its cases. Out of the cases sequenced, 20 out of 65 cases contained TP53 mutations. For cases with TP53 mutations, the upper quartile survival time was 30.22 months. However, for cases without TP53 mutations, an estimated 93.5% of the people were still alive by the end of the study in 136.96 months and thus even an upper quartile survival time could not be calculated. The third was uterine endometrial carcinoma, with 27.8% of its 248 cases containing TP53 mutations. The upper quartile survival time was 36.37 months for cases with TP53 mutations, which was 83.91 months for cases without TP53 mutations. The fourth was thymoma, with 3.3% of its 123 cases containing TP3 mutations. Although this was a small percentage, the survivability of a patient was drastically decreased by the presence of a TP53 mutation. For cancers with TP53 mutations, the median survival time was 25.46 months, and the upper quartile survival time was 12.45 months. In cases without TP53 mutations, the upper quartile survival time was 114.59 months.

Something to be noticed is that in the majority of the cancers analyzed, there were almost no differences in survival for cases with and without TP53 mutations during months 1–10. Only four of the ten cancers that had statistically significant differences in survival had notable differences in survival during months 1–10. In all four cancers, the survival probability was lower for cases with TP53 mutations than those without TP53 mutations. The first was lung adenocarcinoma, where the difference in survival probability between those with and without TP53 mutations (∼10%) stayed around the same throughout the whole Kaplan-Meier estimate. By the tenth month, the difference in survival probability was about 7%. The second cancer was acute myeloid leukemia, where by the tenth month there was about a 45% difference in survival probability. The third was kidney RCC, in which the survival probability difference was around 40% by the tenth month. Lastly, in papillary RCC, the difference in survival probability was around 30% by the tenth month. For the rest of the 10 cancers, there were almost no differences during months 1–10.

### Disease-free survival

Out of the 33 cancers studied, six were found to have statistically significant differences in disease-free survival time. In ovarian serous cystadenocarcinoma, the presence of TP53 mutations was not only beneficial to overall survival, but also to disease-free survival time. The median disease-free survival time was 16.69 months in cases with TP53 mutations and 10.94 months in cases without TP53 mutations. Those with mutations remained disease-free for 1.53 times longer. All cases without TP53 mutations relapsed within 38.41 months, while about 11.05% of those with TP53 mutations remained disease-free until the end of the study at 180.04 months (Fig 6A).

**Fig 6.**
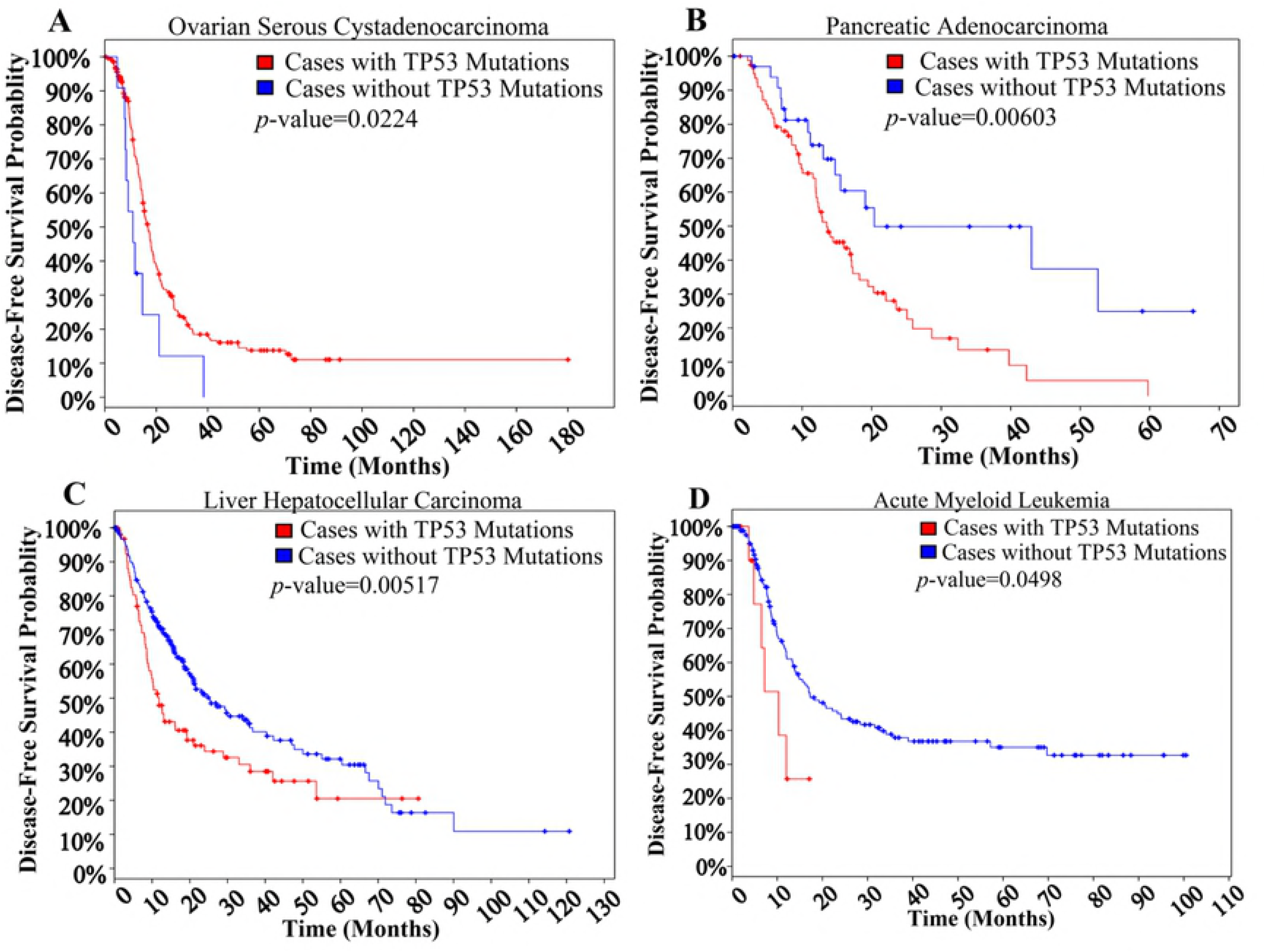
Disease-free survival Kaplan-Meier estimates of select cancers. Data from TCGA studies compiled in cBioPortal for Cancer Genomics.

In pancreatic adenocarcinoma, liver hepatocellular carcinoma, chromophobe RCC, acute myeloid leukemia, and thymoma, TP53 mutations lead to earlier relapse. In pancreatic adenocarcinoma, 69.3% out of 150 cases were found to have TP53 mutations. The median disease-free survival time was 13.53 months in cases with TP53 mutations, and 20.37 months in cases without TP53 mutations. Those without TP53 mutations remained disease-free for 1.51 times longer. However, longer time before relapse did not indicate longer overall survival because although the difference in the time of disease-free survival among those with and without TP53 mutations was statistically significant, there was no statistically significant difference in overall survival time in pancreatic adenocarcinomas. Therefore, cases with TP53 mutations may require more frequent screening and treatment than cases without TP53 mutations due to shortened periods between relapses, but overall survival time would be the same (Fig 6B).

In liver hepatocellular carcinoma, the median disease-free survival time was 11.79 months for cases with TP53 mutations and 25.3 months for cases without TP53 mutations. Overall, those without TP53 mutations lived disease-free for 2.15 times longer. According to the Kaplan-Meier estimates, 20.5% of cases with mutations remained disease-free, while only 10.9% of cases without mutations remained disease-free at the end of the study (Fig 6C).

In acute myeloid leukemia, the median disease-free survival time was 10.3 months for cases with TP53 mutations, and 17.3 months for cases without TP53 mutations. Those without mutations were disease-free for 1.68 times longer than those with mutations (Fig 6D).

Chromophobe RCC tended to not relapse in cases without TP53 mutations. In the first 26.84 months, while 32.1% of cases with TP53 mutations relapsed, 100% of cases without TP53 mutations remained disease-free. After 65.34 months, 32.1% of cases with TP53 mutations relapsed, while 92.2% of cases without TP53 mutations remained disease-free.

In thymoma, although TP53 mutations were less frequent than in other tumors, having TP53 mutations significantly decreased time before relapse. The median disease-free survival time was 9.72 months and the upper quartile time was 7.36 months in cases with TP53 mutations. For cases without TP53 mutations, no median disease-free survival time could be calculated since <50% of cases relapsed. The upper quartile time for cases without TP53 mutations was 115.05 months, which was greater than the median time for cases with TP53 mutations. In addition, the upper quartile value for cases without TP53 mutations was 15.63 times that of cases with TP53 mutations. Therefore, the presence of TP53 mutations was very detrimental in thymoma cases.

### Specific TP53 mutations as prognostic markers

For some cancers, specific TP53 mutations can be used as prognostic markers. This can be done in cancers with a large number of samples containing TP53 mutations. One Kaplan-Meier estimate was generated for liver hepatocellular carcinomas, showing the effect of the R249S mutation (the most common p53 mutation in liver hepatocellular carcinoma) on survival compared to cases with and without TP53 mutations. The median survival time for cases with the p53 R249S mutation was 11.30 months compared to 45.07 months for cases with TP53 mutations (including R249S) and 60.84 months for cases without TP53 mutations. Since the differences in survival time among the three groups was statistically significant, this specific mutation is an indicator for poor prognosis in liver hepatocellular carcinoma (Fig 7).

**Fig 7.**
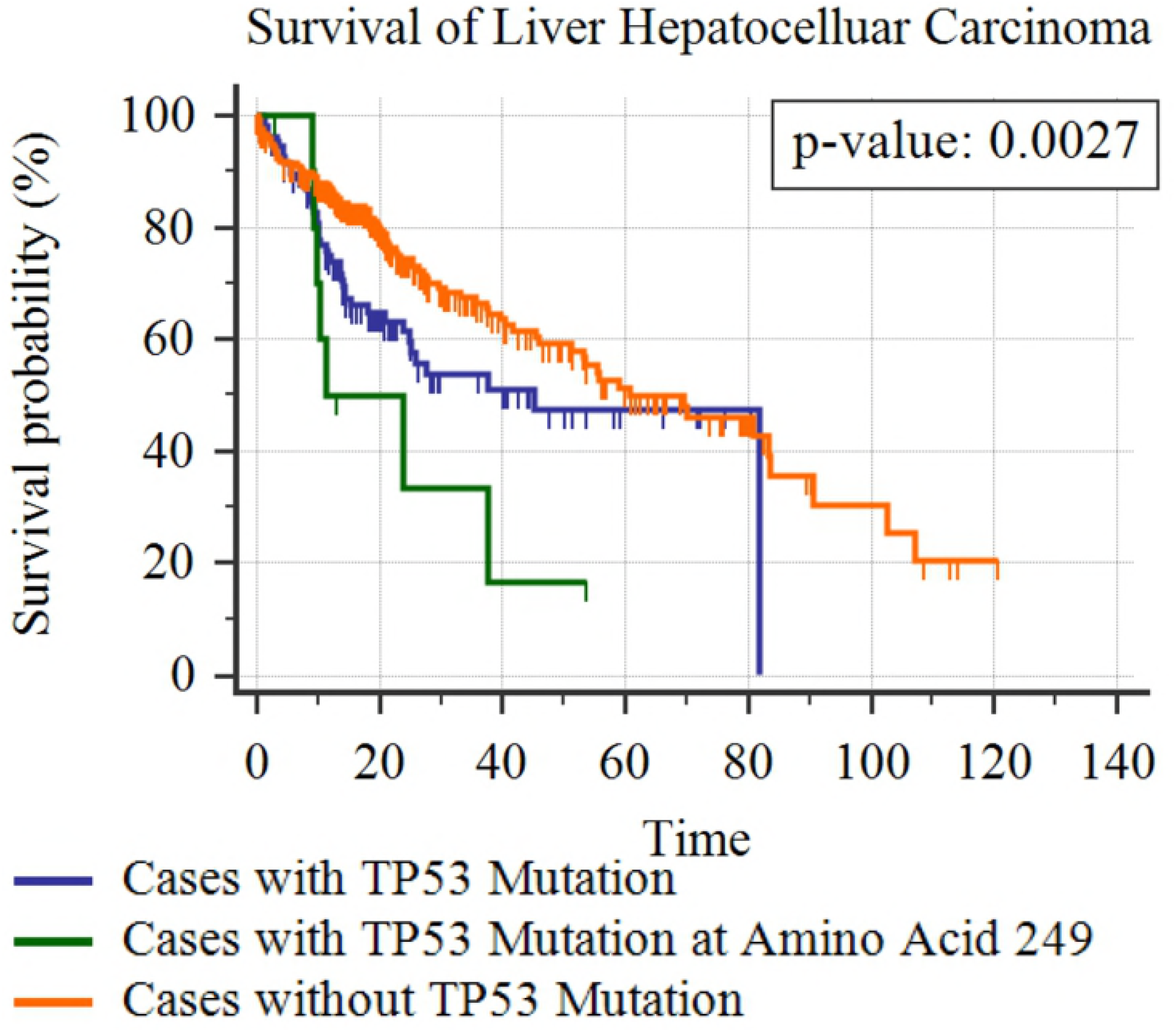
Survival Kaplan-Meier estimate of liver hepatocellular carcinoma cases with p53 R249S mutation. Data from TCGA studies compiled in cBioPortal for Cancer Genomics.

## Discussion

This study used TP53 mutations instead of immunohistochemistry (IHC) data even though both were available. Since the p53 protein sometimes accumulates when there is no mutation and sometimes does not accumulate when there is a mutation present, the presence of false-positives and false-negatives making IHC less reliable than sequencing for TP53 mutations [15]. Older research uses IHC, but due to the availability of faster and cheaper modern sequencing methods, more genetic data on TP53 has become available for analysis.

Some mutations affected the survival time or time before relapse, while others had no effect. In 26 of the 33 cancers studied, there was data on survival time. Ten cancers (ovarian serous cystadenocarcinoma, lung adenocarcinoma, liver hepatocellular carcinoma, head and neck squamous cell carcinoma, acute myeloid leukemia, clear cell RCC, papillary RCC, chromophobe RCC, uterine endometrial carcinoma, and thymoma) had statistically significant differences in survival time between patients with and without TP53 mutations. Of the 33 cancers studied, there were 21 cancers with relapse data, and six cancers (ovarian serous cystadenocarcinoma, pancreatic adenocarcinoma, liver hepatocellular carcinoma, chromophobe RCC, acute myeloid leukemia, and thymoma) had statistically significant differences in disease-free survival time between patients with and without TP53 mutations.

With these data, TP53 mutations can be predictive markers as to how long patients will survive, and when the cancer may relapse. For example, with ovarian serous cystadenocarcinomas, one can predict that if a patient does not have a TP53 mutation, then she will have a 45% chance of surviving to month 31, while another patient with the same disease will have a 70% chance of surviving to month 31 if she has a TP53 mutation (Fig 5A). This adds accuracy to preexisting survival estimates that do not consider TP53 mutations as a factor. The same applies to disease-free survival time. In ovarian cystadenocarcinomas, while 55% of patients without TP53 mutations are expected to relapse by month 11, only 30% of patients with TP53 mutations are expected to relapse in the same amount of time (Fig 6A).

Past studies have determined the function of the p53 protein and studied the development TP53 mutations. Since the establishment of the IARC TP53 Database in 1991, many researchers have analyzed and used its data to identify the different TP53 polymorphisms that exist in human populations [16]. Researchers have also identified the causes of TP53 mutations, including geographic differences and known carcinogens [11]. In addition, the effects of TP53 mutations on the function and structure of the p53 protein were studied, which led to the identification of the functional domains of p53 in 2006 [3, 17].

Using TP53 mutation data from the newest IARC Database, which is the April 2016, R18 release, and data on cBioPortal, which compiles all of the data from the most recent TCGA cancer studies, this study is the first to represent the newest data available. Applying previous knowledge of p53 functional domains and their locations, we were able to identify the distribution of mutations at exons/introns to determine that the DBD, which is responsible for binding p53 to DNA to regulate cell growth, was affected the most by TP53 mutations. By separating mutations into somatic and germline mutations, we found that germline mutations were common at introns and in the TD, AD1, and AD2, which was rare among somatic mutations.

Using survival and genomic data in multiple TCGA studies on cancers, this study compared Kaplan-Meier estimates of cases with and without TP53 mutations to predict survival time and disease-free survival time. Unlike preexisting studies, we examined the prognostic effect of TP53 mutations in a whole variety of cancers, instead of one specific cancer, and found that survival was significantly affected in 10 different kinds of cancers. This study is also one of the only studies that looked at the significance of TP53 mutations on the time before relapse. We also discovered that TP53 mutations in ovarian serous cystadenocarcinomas, which mostly occurred in the DBD, increased the survival and the disease-free survival time of patients, contrary to a study published in 2007 that said “TP53 mutations within the [DBD] have been repeatedly associated with shorter survival” [18]. The distribution of TP53 mutations was also used to identify the most prevalent mutations in certain cancers. The mutations in liver hepatocellular carcinoma were analyzed and the most common one was R249S. Survival data for cases with this specific mutation showed that it negatively affected the prognosis of patients significantly. Due to a previous lack of data, specific TP53 mutations were not considered to be utilized as prognostic markers until now.

Nevertheless, for some of the cancers, more data is necessary. For seven of the cancers, survival data was not available, and for twelve of the cancers, disease-free survival time data was not available either. Also, for some of the cancers, only small amounts of data were available since only a small percentage of cases had TP53 mutations. Lastly, some Kaplan-Meier estimates could be misleading because of censoring that results in horizontal and vertical lines which indicate loss of data [19]. More data can lead to better accuracy for cancers like clear cell RCC, where there were only nine cases of TP53 mutations and four of them were censored.

## Methods

Data was downloaded from the International Agency for Research on Cancer (IARC) TP53 database on all 28,000+ somatic and 800+ germline mutations [2]. Python was then used to scan through the IARC data to generate graphs via Matplotlib, an open source plotting library that can be used in IPython (Jupyter Notebook), and Glue, a tool built on top of the standard scientific libraries of Python. Exon/intron distribution maps, along with codon distribution maps were generated for both somatic and germline mutations. The mutational hotspots were also located, and mutational hotspots were analyzed for germline mutations to see if families with LFS or LFL had mutations at certain codons. The image of the domains of the p53 protein was drawn in Google drawings using data from “The Functional Domains in p53 Family Proteins Exhibit Both Common and Distinct Properties” [3].

Genomic data from various cancer studies can be found in cBioPortal for Cancer Genomics, available online at cbioportal.org [9, 10]. The data sets of the most recent The Cancer Genome Atlas (TCGA) cancer studies were selected in the portal, which resulted in 33 studies and 7,893 cases in total. Then a query was performed for TP53 mutations, from which cBioPortal generated a summary of the percentage of cases with TP53 mutations for each cancer. Of these studies, survival data and overall survival Kaplan-Meier estimates were downloaded from cBioPortal, and patients with and without TP53 mutations were compared. This data was analyzed to obtain the total number of cases, the number of cases deceased, and the survival times of patients with and without TP53 mutations for individual cancers. The statistical significance of differences in survival time was determined by logrank *p*-values. A value of 0.05 or less was considered statistically significant.

Studies with data on relapse and disease-free time were also selected. Disease-free survival Kaplan-Meier estimates were downloaded and analyzed. Estimates with logrank *p-*values of 0.05 or less were considered statistically significant. Disease-free survival data included the total number of cases, number of cases relapsed, relapse time of all the patients, censored data, and whether a TP53 mutation was present. The data was also compared with the survival data to determine whether relapse affected overall survival.

Lastly, data was selected for specific TP53 mutations in liver hepatocellular carcinoma cases. Survival data of cases with the most common mutation (R249S) in this cancer were used to generate a Kaplan-Meier estimate using MedCalc, and the survival time of cases with this specific mutation, cases with TP53 mutations, and cases without TP53 mutations were compared.

## Conclusion

The presence of TP53 mutations resulted in poor prognosis in 9 cancers (lung adenocarcinoma, liver hepatocellular carcinoma, head and neck squamous cell carcinoma, acute myeloid leukemia, clear cell RCC, papillary RCC, chromophobe RCC, uterine endometrial carcinoma, and thymoma). Only in ovarian serous cystadenocarcinoma did the presence of TP53 mutations lead to good prognosis. With current data, TP53 mutations can be reliable prognostic markers on their own. These results are supported with current research on TP53 and its clinical value [20]. However, when combined with other factors, the accuracy of survival and relapse estimates can be further improved. The most common p53 mutation (R249S) in liver hepatocellular carcinomas was found to be an indicator for poor prognosis.

In future studies, when more data becomes available, by identifying and analyzing the most common TP53 mutations and the cancers they are prevalent in, it will be possible to predict the invasiveness or aggressiveness of cancers based upon whether they contain TP53 mutations or not, and in cases with TP53 mutations, based upon which specific TP53 mutations are present [21]. This information can be useful in determining the progression of cancers, which is crucial when deciding the type of treatment to be used. Also, the effects of specific mutations in cancers can be investigated to a deeper extent on the molecular level to improve accuracy of TP53 mutations as prognostic markers. This study demonstrated that TP53 mutations have statistically significant effects on survival and disease-free survival time, and thus can be successfully used as prognostic markers.

Specific germline TP53 mutations and the cancers that they are associated with can be identified to predict the cancers certain individuals and their children are predisposed to, which can help implement screening procedures that allow for earlier detection and treatment. Individuals with a certain germline TP53 mutation can be screened for specific cancers that they are at risk of developing, thus increasing the likelihood of finding and successfully treating premalignant tumors and growths through preventative care. However, more data on germline TP53 mutations is required to establish reliable criteria for screening. As of now, there are still less than one thousand samples in the IARC TP53 Database, the largest database in the field. Nevertheless, with increasing data availability, future studies may yield further insights.

